# Thermodynamics of droplets undergoing liquid-liquid phase separation

**DOI:** 10.1101/2021.04.01.438092

**Authors:** Subhadip Biswas, Biswaroop Mukherjee, Buddhapriya Chakrabarti

## Abstract

We study the thermodynamics of binary mixtures wherein the volume fraction of the minority component is less than the amount required to form a flat interface. Based on an explicit microscopic mean field theory, we show that the surface tension dominated equilibrium phase of a polymer mixture forms a single macroscopic droplet. A combination of elastic interactions that renormalize the surface tension, and arrests phase separation for a gel-polymer mixture, stabilize a micro-droplet phase. We compute the droplet size as a function of the interfacial tension, Flory parameter, and elastic moduli of the gel. Our results illustrate the importance of the rheological properties of the solvent in dictating the thermodynamic phase behavior of biopolymers undergoing liquid-liquid phase separation.

---

Membraneless compartmentalisation in cells that are driven by phase-separation processes due to changes in temperature or *pH*, and maintained by a non-vanishing interfacial tension, is one of the most exciting recent biological discoveries [1–4]. These membraneless compartments composed bio-molecular condensates have been implicated in important biological processes such as transcriptional regulation [5], chromosome organisation [6] and in several human pathologies *e*.*g*. Huntington’s, ALS *etc*. [7]. Self-assembly processes that lead to organelle formation however need to be tightly regulated such that the phase separated droplets do not grow without bound and remain small compared to the cell size. Understanding the regulatory processes that controls droplet size in cellular environments is therefore a crucial interdisciplinary question. The two candidate mechanisms proposed for arresting droplet growth are (i) incorporation of active forces that break detailed balance [8, 9], and (ii) non-equilibrium reaction mechanisms which couple to the local density field [3]. Although biological systems are inherently out of equilibrium, an estimation of diffusion constant of bio-molecules indicates that non-equilibrium effects are negligible on length-scales beyond microns and timescales beyond microseconds. Hence, the framework of equilibrium thermodynamics can be readily applied to analyse biological phase separation in cells.

For synthetic polymer mixtures, in the absence of active processes, droplet growth is limited by the elastic interactions of the background matrix that alters the thermodynamics of phase separation [10, 11]. Recent experiments on mixtures of liquid PDMS and flourinated oil in a matrix of cross-linked PDMS show the dependence of the droplet size on the nucleation temperature and the network stiffness [12, 13]. Despite theoretical attempts [14, 15] a complete understanding of elasticity mediated arrested droplet growth is still lacking.

The connection between coarsening phenomena and network elasticity is an important, and exciting area of research across several disciplines, biological regulation of cellular function [1–4], tailoring mechanical properties of materials [16–18], controlling morphology [19–21], size of precipitates in food products [22, 23], and even growth of methane bubbles in aquatic sediments [24–26].

In this letter, we develop a consistent thermodynamic formalism to compute the equilibrium radius of the droplet of the minority phase in (a) binary polymer, and *(b)* a polymer-gel mixture, using mean-field theories utilising the Flory-Huggins [27] and the Flory-Rehner [28] free energies, respectively. A parallel tangent construction for droplets, used to obtain the densities of coexisting phases is presented. This procedure is a generalisation of the common tangent construction for flat interfaces and in the thermodynamic limit allows us to compute the equilibrium radius of a single droplet. For phase separation processes in mixtures with a gel component, elastic interactions limit droplet growth stabilising a phase with multiple droplets, in the correct parameter regime [29].

The thermodynamic formalism to understand phase separation is as follows: an unstable mixture of composition *ϕ*_0_ splits into two coexisting phases in a slab-like geometry respecting volume and mass conservation, with the equilibrium configuration being a minimum of the free energy (Fig.(1)(a)). The densities of the two coexisting phases are *ϕ*_*in*_, and *ϕ*_*out*_ respectively, with *V*_*d*_ denoting the volume occupied by phase with density *ϕ*_*in*_, and ℱ_*b*_(*ϕ*) is the Helmholtz free-energy per unit volume (in units of 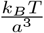). The volume fraction of the solvent with density *ϕ*_*in*_, is 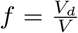, and the free-energy density ℱ (*ϕ*) of the planar configuration (Fig.(1)(a)) is given by,

**FIG. 1.**
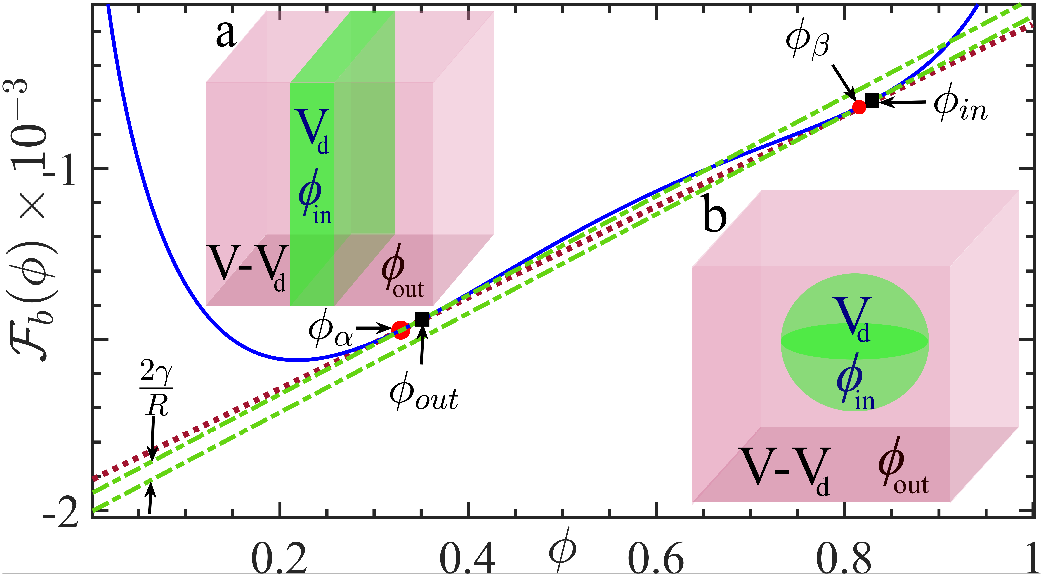
A (i) common tangent (solid pink line) and a (ii) parallel tangent (dashed green line) construction for planar interfaces (a) and droplets (b) (with volume *V*_*d*_) for a binary polymer mixture obtained using a Flory-Huggins functional with *χ* = 1.2*χ*_*c*_, *N*_*A*_ = 100, *N*_*B*_ = 200. Coexistence densities inside *ϕ*_*in*_ and outside *ϕ*_*out*_ droplet approach the values obtained for a flat interface *ϕ*_*α*_, and *ϕ*_*β*_ in the thermodynamic limit.

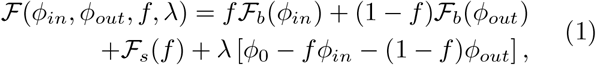

where ℱ_*s*_ = 2*γV* ^*−*1*/*3^, corresponds to the surface energy with *γ* being the surface tension, *V* the volume of the system considered, and *λ* a Lagrange multiplier that enforces the mass conservation constraint.

A calculation of the equilibrium thermodynamics proceeds via minimising the free energy in Eq.(3) w.r.t to the independent quantities *ϕ*_*in*_, *ϕ*_*out*_, *f*, and *λ*. The constrained minimization of the free-energy function in Eq.(1) w.r.t. *ϕ*_*in*_, *ϕ*_*out*_ and *f* leads to the common tangent construction

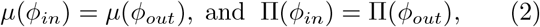

where *µ*(*ϕ*), and Π(*ϕ*) refers to the chemical potential and the osmotic pressure of the phases respectively. Eq.(2) ensures chemical, and mechanical equilibrium (see SI). Thermal equilibrium is ensured as calculations are carried out in a constant temperature ensemble. We obtain coexistence densities *ϕ*_*in*_ and *ϕ*_*out*_ from Eq.(2). The solvent fraction *f* is obtained by minimising the functional w.r.t *λ, i*.*e ∂*ℱ(*ϕ*_*in*_,*ϕ*_*out*_, *f, λ*)*/∂λ* = 0, which yields, 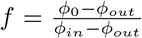 For a planar interface, the surface energy term does not explicitly depend on the solvent volume fraction *f*. Consequently, the minimisation conditions lead to four uncoupled equations (SI) and a knowledge of the coexistence densities *ϕ*_*in*_ and *ϕ*_*out*_ is enough to determine *f*. As evident from Eq.(1), the effect of the surface energy term vanishes in the thermodynamic limit, *i*.*e*., as volume *V*→ ∞. In contrast, a spherical droplet geometry introduces a non-trivial coupling among the minimisation conditions and a knowledge of the volume, *V*, of the system is required to obtain the equilibrium configuration.

Spherical droplets of the minority phase arise in finite systems when the volume fraction is less than a critical value [30, 31]. The thermodynamics in such situations differ from the common tangent construction and leads to the classical Gibbs-Thomson relations [3]. Fig.(1)(b) shows an unstable system of density *ϕ*_0_, that phase separates into a background matrix of density *ϕ*_*out*_ and a single droplet of radius *R* of density *ϕ*_*in*_ in a finite box of volume *V*. Assuming an ansatz of a phase separated mixture comprising of *N* spherical droplets of identical radius *R*, (referred to as the micro-droplet phase henceforth), the solvent volume fraction is given by 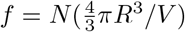.The free energy of the micro-droplet phase is therefore ℱ= *f* ℱ_*b*_(*ϕ*_*in*_)+(1 − *f*)ℱ_*b*_(*ϕ*_*out*_) + ℱ_*s*_(*f*), where 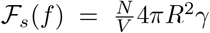, accounts for the interfacial energy between the droplet and the background phase. By imposing the mass conservation constraint and expressing the surface energy term in terms of the solvent fraction *f*, the free energy per unit volume is given by

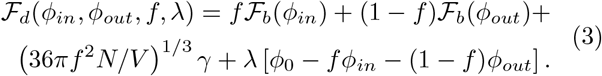

The surface energy of the droplet depends on the solvent fraction *f* on account of the its spherical shape. The equilibrium conditions therefore lead to four coupled equations, involving the yet unknown system volume *V*. The chemical and mechanical equilibrium conditions for the micro-droplet phase involving the co-existing densities translates to, *µ*(*ϕ*_*in*_) = *µ*(*ϕ*_*out*_) and 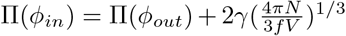, where the extra term in the pressure equation accounts for the Laplace pressure acting across the interface. We carry out a minimisation procedure akin to the planar interface to obtain the solvent volume fraction *f*, and the coexistence densities inside and outside the droplet, *ϕ*_*in*_ and *ϕ*_*out*_ respectively for a given box volume *V*. In the absence of elastic interactions the equilibrium phase corresponds to a single droplet of the minority phase, i.e. *N* = 1 in Eq.(3). The radius of the drop is determined in terms of the coexistence densities and is given by

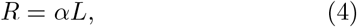

Where 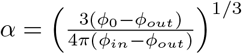, and *L* = *V* ^1*/*3^ is the length of the cubic box. We apply the framework to compute the radius of the minority phase droplet of a binary polymer mixture described by a Flory-Huggins free energy in the thermodynamic limit *i*.*e. V*→ ∞, performing our calculation for different box volumes *V*. The surface tension *γ* for the micro-droplet phase is taken to be the same as that of a planar interface.

The thermodynamics of binary polymer mixtures is well described by the Flory-Huggins free-energy 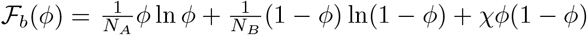, where *N*_*A*_, and *N*_*B*_ are the lengths of *A* and *B* polymers respectively, and *χ* is the mixing parameter. For *χ > χ*_*c*_, where *χ*_*c*_ is the value of the mixing parameter at criticality, the mixture is unstable and spontaneously phase separates into low and high density phases determined by the minimisation conditions. We consider an unstable polymer mixture with *N*_*A*_ = 100, *N*_*B*_ = 200, and an initial composition *ϕ*_0_ = 0.35, and *χ* = 1.2*χ*_*c*_. Fig.(1)(a) shows the common tangent construction which yields the coexistence densities *ϕ*_*α*_ = 0.235 and *ϕ*_*β*_ = 0.885 for a flat interface. If the amount of material is not enough, the minority phase forms a droplet whose coexistence densities outside *ϕ*_*out*_ and inside *ϕ*_*in*_ are determined by the parallel tangent construction (Fig.(1)(b)) as a function of the box volume *V*. A combination of the parameters *ϕ*_0_, *ϕ*_*α*_ and *ϕ*_*β*_ determines that the volume fraction of the solvent phase *f* ≈ 0.17. The equilibrium phase is a single drop. To obtain the coexistence densities and the droplet radius in the thermodynamic limit, we perform parallel tangent constructions for cubic boxes of lengths *L* = 160,… 10^4^ using Eq.(4).

Fig.(2) shows a finite size scaling analysis of the droplet radius *R* in units of the box-size *L*, (*R/L*) as a function of 1*/L*. The thermodynamic limit 1*/L* → 0 corresponds to the *y*-intercept *R/L* ≈0.34 for the Flory parameters listed above. The numerical derivative of *R/L* w.r.t. *L* approaches zero in this limit (Fig.(2) inset). The solvent fraction *f*, is also a function of the systems size (*f* ∼ (*R/L*)^3^). The coexistence densities calculated from Eq.(4) are functions of *L* and can be quantified in terms of their deviation from the coexistence densities for a planar interface, *i*.*e*.,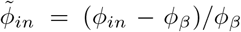 and 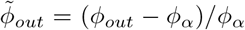 As shown in Fig.(2) *ϕ*_*out*_→ *ϕ*_*α*_, and *ϕ*_*in*_→ *ϕ*_*β*_ in the thermodynamic limit.

**FIG. 2.**
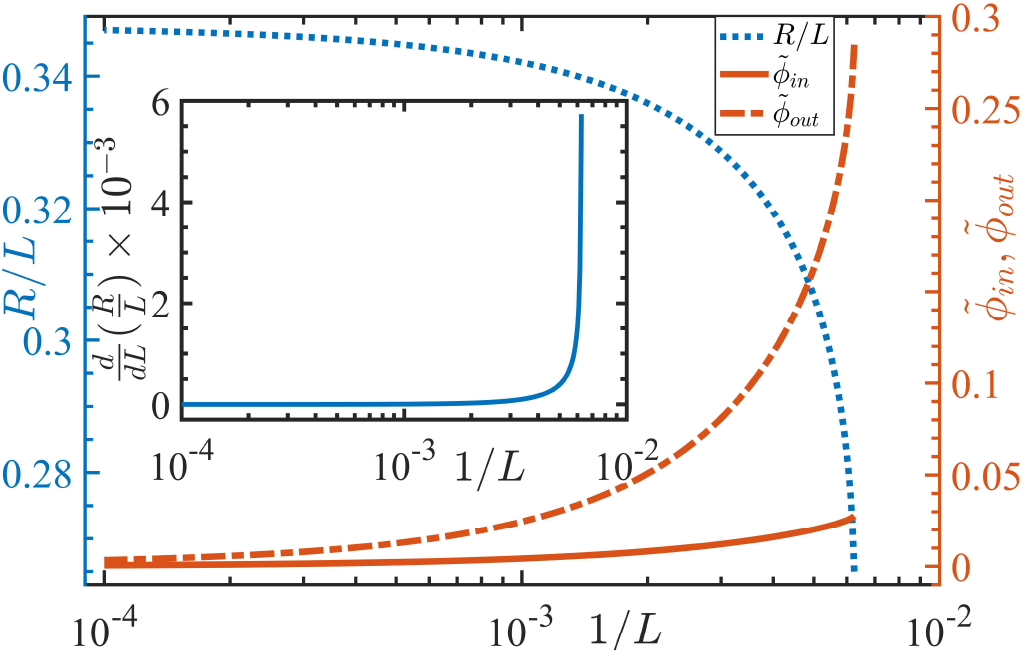
Finite size scaling of equilibrium drop radius *R*(*L*)*/L*, of a phase separated binary polymer mixture. Coexistence densities inside and outside the droplet 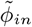 and 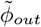 approaches the coexistence values obtained from a common tangent construction as *L →* ∞. Inset shows the rate of change of the radius approaches zero as *L →* ∞.

The Helmholtz free-energy per unit volume of the micro-droplet configuration of a gel-solvent mixture, with *N* droplets (see Fig.(4) inset), is given by

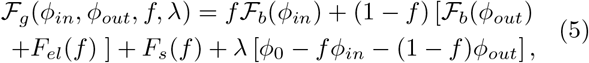

where ℱ_*b*_(*ϕ*) is the Flory-Huggins free energy in the limit *N*_*B*_→ ∞ owing to the macroscopic gel size. Thus, ℱ_*b*_(*ϕ*) = *ϕ* ln *ϕ* + *χϕ*(1 −*ϕ*), and the mixing parameter, 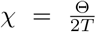, with Θ = 1183*K*. We perform thermodynamic calculations of the gel-solvent mixture at *T* = 350*K*. The surface-energy per unit volume in Eq.(5), 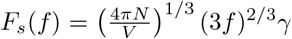 is expressed in terms of the solvent fraction *f* using the relation between the drop radius and the number density, *i*.*e*., 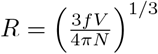. The elastic part of the free-energy density is also a function of the fraction, *f*, (see SI) and is given by

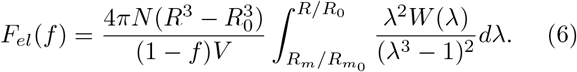

To incorporate the effects of the finite stretch-ability of the gel, we adopt the Gent model [32, 33]. The elastic free energy density has the form, 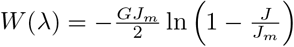, where 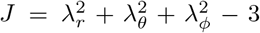, with *λ*’s corresponding to the strains in the radial, azimuthal, and polar directions, *J*_*m*_ ∼ 10^6^ is the stretching limit of the network, and *G* is the shear modulus. The shear modulus is related to the microscopic parameters via the relation, 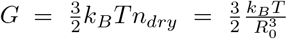, where *n* _*dry*_ and *R*_0_ are the average cross-link density and the mesh size of the dry gel respectively[34] (see SI). Due to the volume-preserving nature of the deformation, *λ*_*r*_ = 1*/λ*^2^ and *λ*_*ϕ*_ = *λ*_*θ*_ = *λ* and its magnitude is bounded, *i*.*e*., 0 *< J/J*_*m*_ *<* 1 [32]. The energy minimisation conditions w.r.t the independent variables as outlined earlier, leads to a modified chemical and mechanical equilibrium conditions: *µ*(*ϕ*_*in*_) = *µ*(*ϕ*_*out*_) and 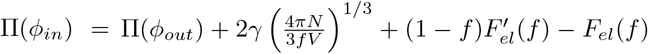. These conditions lead to a set of coupled equations that we solve numerically to yield the four unknown variables, *ϕ*_*in*_,*ϕ*_*out*_,*f*, and *λ*, associated with each droplet number, *N*. Due to the functional form of ℱ_*b*_(*ϕ*) in Eq.(5), where the entropic term associated with the gel is absent, *ϕ*_*in*_ is located at the unstable part of the free-energy surface. Despite this, the micro-droplet phase is in mechanical equilibrium due to the elastic energy residing in the gel-matrix.

We substitute the equilibrium values of the coexistence densities and solvent fraction into the original free-energy expression in Eq. (5), to obtain a free energy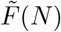, as a function of the number of droplets *N*. The minimisation of 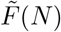 w.r.t *N* yields *N*_*m*_, the optimal number of droplets of the micro-droplet phase.

Fig.(3)(a) shows the free-energy 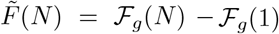 (Eq.(5)) as a function of the number of droplets, once the coexistence densities have been obtained for a cubic box of side *L* = 200 and the surface tension *γ* = 1.67 ×10^*−*3^ (in units of *k*_*B*_*T/a*^2^). It is evident that this is a convex function, with a well defined minimum occurs around *N*_*m*_≈ 30. The inset shows the contrasting behaviour of 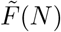 for a binary polymer mixture. In the absence of elastic interactions, surface tension dominates the thermodynamics and a phase with a single droplet is the equilibrium state corresponding to the free energy minimum. The convex nature of the free energy 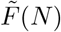 arises from a balance between the surface, elastic, and bulk free energies of the micro-droplet phase. As the number of droplets *N* increases, the surface energy monotonically increases on account of the increase of the total interfacial area. In contrast, the elastic energy monotonically decreases as a function of *N*, since an increase in the number of droplets translates to smaller sized drops and less deformation of the gel matrix. The elastic free energy has a lower bound corresponding to a minimum droplet of size *R/L* ∼ *a*, length of a monomer. The combined effect of these two contributions to the free energy therefore stabilizes the micro-droplet phase. The bulk free energy is nearly independent of *N*. Fig.(3)(b) shows the variation of the different components of the total free energy as a function of the number of droplets *N*.

**FIG. 3.**
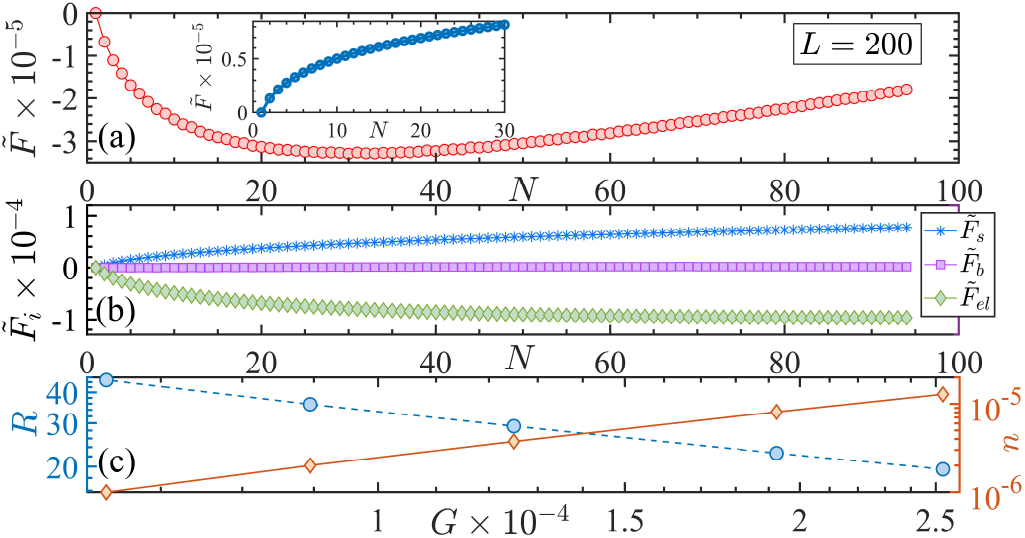
Total free energy, (a), as a function of the number of droplets *N* showing a minimum at *N*_*m*_ ≈ 30 for *L* = 200. Surface energy *F*_*s*_, increases, the elastic energy *F*_*el*_, decreases, whereas, the bulk free energy ℱ_*b*_(*ϕ*)is almost independent of *N* as shown in (b). Panel (c) shows the dependence of the droplet radius *R* and the number density of the droplets *n* and the shear modulus of the gel *G*.

Fig.(3)(c) shows the variation of number density *n* = *N*_*m*_*/V*, and droplet radius *R* as a function of the shear modulus *G*. The shear modulus *G* is tuned by varying the mesh size, *R*_0_, of the gel. We compute the number density by minimizing 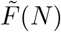 w.r.t. *N* and determine the drop radius using 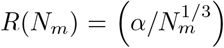 *L* for a given shear modulus *G*. As shown, the radii of the droplets decrease (and hence the number density increases commensurately) as the gel becomes stiffer. Our results corroborate the recent experimental findings on liquid-liquid phase separation in elastic networks [12, 13]. The convex nature of 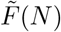 as a function of *N* is independent of the system size *L* as shown in Fig.(4)(a)). Fig.(4)(c) shows that the equilibrium number density of droplets *n* = *N*_*m*_*/V*, and the droplet radius *R*(*N*_*m*_) have reached a thermodynamic limit and are independent of the system size *L*. Fig.(4)(b) shows the dependence of 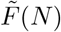 as a function of the surface tension, *γ* as it is varied between 0.0025, and 0.004 while keeping the shear modulus of the gel solvent mixture fixed at *G* = 0.19 × 10^*−*3^ (in units of *k*_*B*_*T/a*^3^). The free energy minimum shifts to smaller values of *N*_*m*_ with increasing surface tension as shown in Fig.(4)(b). Below a critical value *γ*_*c*_ ≈3.45× 10^*−*3^, a micro-droplet phase is the equilibrium configuration, with the number of droplets increasing almost linearly with decreasing surface tension, (*γ*_*c*_−*γ*). A schematic of the micro-droplet phase is shown in Fig.(4)(e). Above *γ*_*c*_ a single macroscopic droplet of the minority component is the thermodynamic equilibrium phase.

**FIG. 4.**
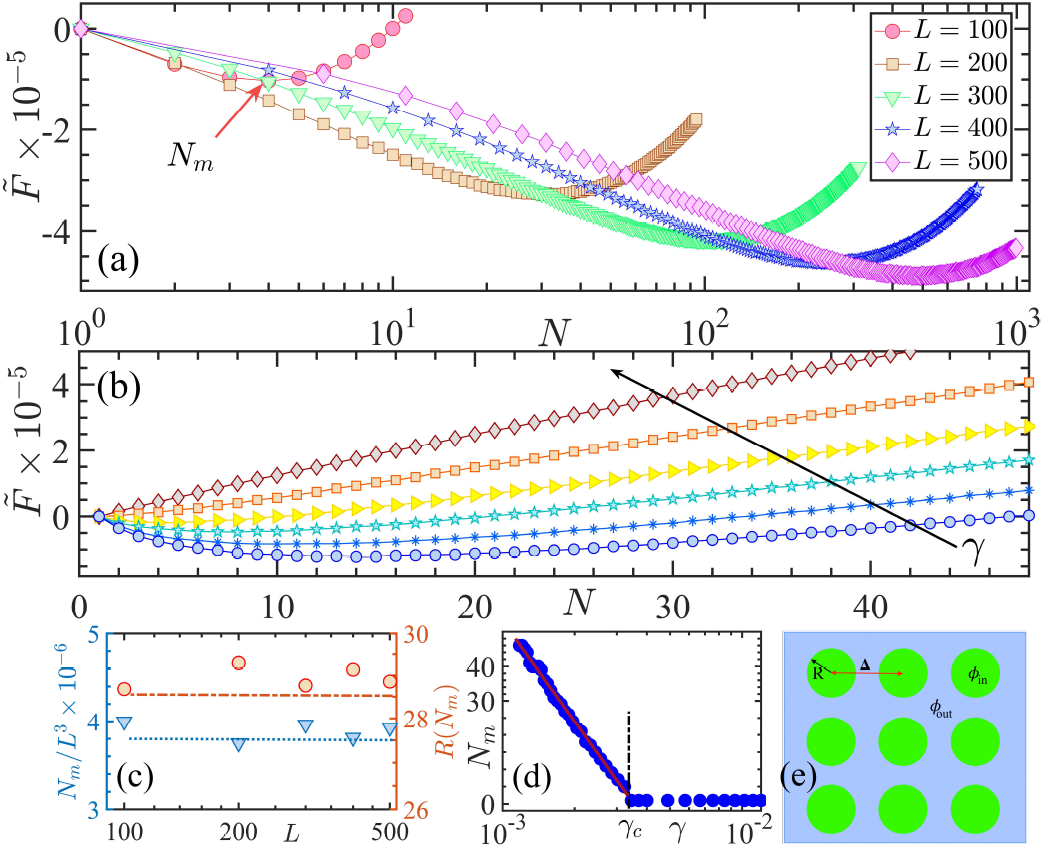
The variation of the free energy of the micro-droplet phase 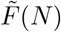 as a function of the number of droplets, *N* for different system sizes *L* = 100, 200,… 500 (panel (a)), and surface tension, *γ* = 0.0025,… 0.004 (b). The shear modulus *G* = 0.19 *×*10^*−*3^*k*_*B*_*T/a*^3^ and box size *L* = 200 is held fixed when the surface tension is varied. The number of stable droplets*N*_*m*_, and their radii *R*(*N*_*m*_) as a function of system size *L* is shown in (c). A single macroscopic drop becomes unstable below a surface tension *γ*_*c*_ (d) and a micro-droplet phase (e) results.

**FIG. 5.**
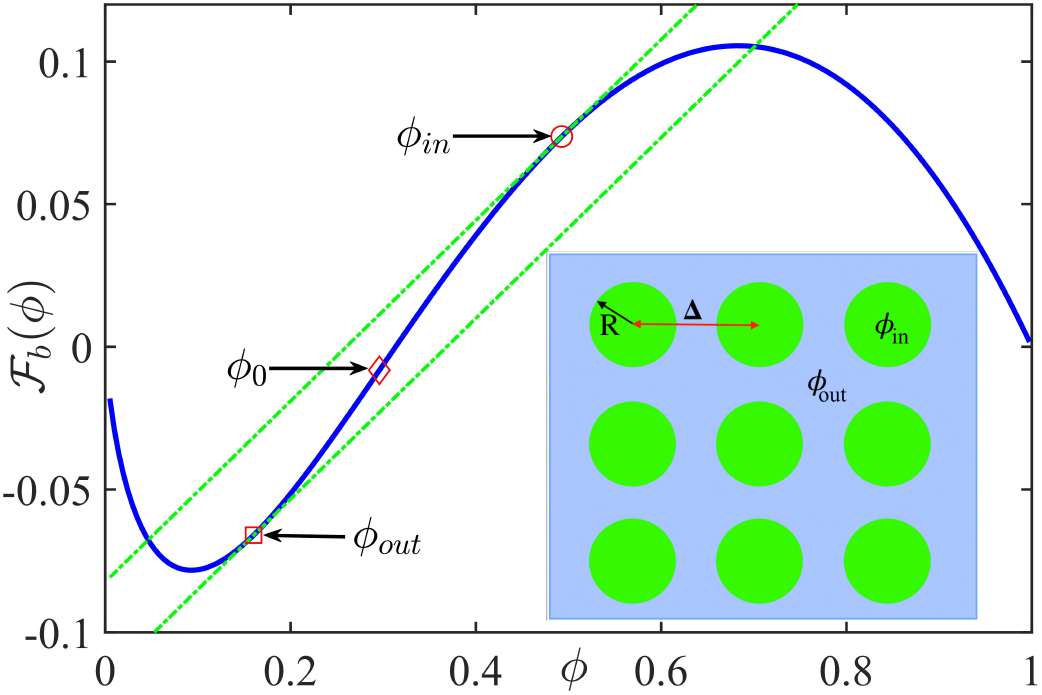
A schematic figure denoting the bulk free energy, ℱ_*b*_(*ϕ*), the set of parallel tangents locating the coexistence densities *ϕ* _*in*_ and *ϕ* _*out*_ for a system with composition *ϕ* _0_.

**FIG. 6.**
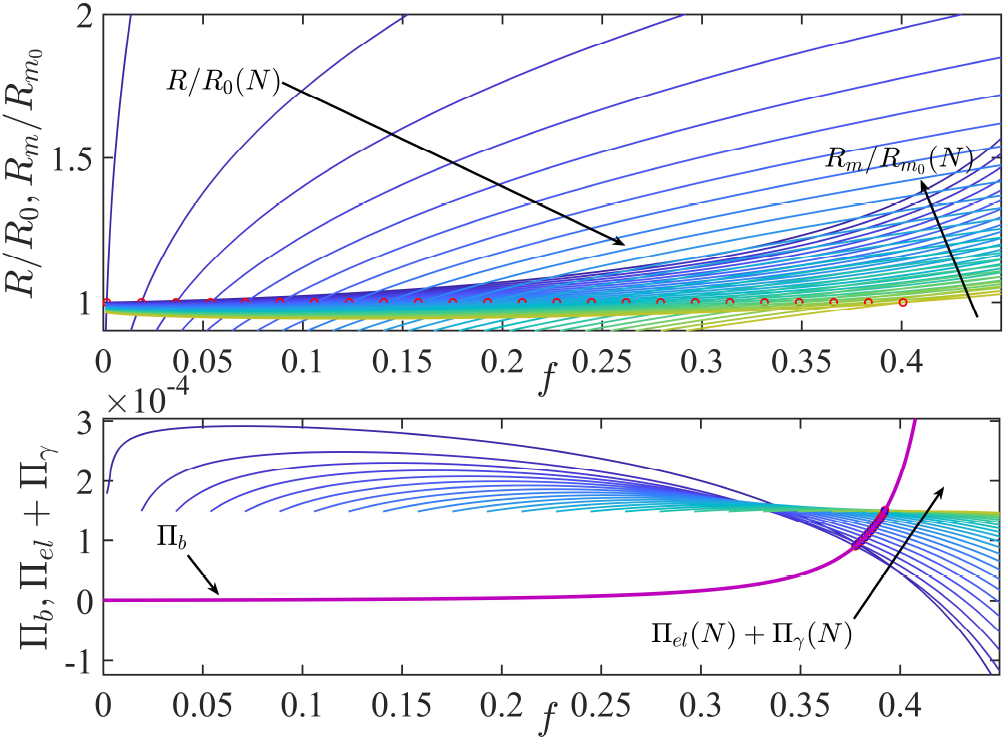
The upper panel shows the upper and lower limits of the integral appearing in the computation of the elastic free-energy, as a function of the solvent fraction *f*, for various number of droplets *N*. The lower panel shows the three terms appearing in the equation for mechanical equilibrium as a function of *f* and their points of intersection, which yields the equilibrium value of *f*. This is again repeated for several number of droplets *N*.

In summary, we consider phase separation in an elastic medium, where the background matrix influences the equilibrium thermodynamics of the mixture. Previous studies consider the background matrix as an inert phase [14, 15]. For composition regimes where the solvent is a minority phase and there is a dearth of material to form a flat interface, solvent-rich droplets coexist with the majority phase. We demonstrate, via explicit meanfield theory calculations, that the dispersed micro-droplet phase is indeed a thermodynamic minimum for a gelsolvent mixture. A competition between surface tension and network elasticity stabilizes this phase. When the surface-tension exceeds a critical value, a single macroscopic droplet is the stable thermodynamic phase. Our framework is generic and can be applied to any system with a bistable free energy [35].

We have considered monodisperse drops as a candidate for the micro-droplet phase. In experiments a certain amount of polydispersity is inevitable. A variational formulation allowing for different ansatz that accounts for polydispersity has not been attempted. Elastically mediated phase transitions admit a third thermodynamic phase, where the gel network partially wets and intrudes the solvent rich droplets[29]. Our formalism may be extended to incorporate such phases. We posit that a model incorporating phase separation and elastic effects is the minimal model to study phase separation in biomolecular condensates.

## ACKNOWLEDGMENTS

SB and BC thank University of Sheffield, IMAGINE: Imaging Life grant for financial support. BC and BM acknowledge funding support from EPSRC via grant EP/P07864/1, and P&G, Akzo-Nobel, and Mondelez Intl. Plc. The authors thank Dr. S Kundu for a critical reading of the manuscript.

## I. APPENDIX

### A. The planar interface

Consider the geometry of the plane interface in the inset (a) of Figure 1 of the main manuscript. The Helmholtz free-energy per unit volume of this system can be written in the following form,

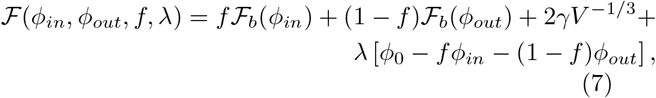

where ℱ _*b*_(*ϕ*) is the free-energy per unit volume of the bulk, *f* is the faction of the solvent phase, *γ* is the surface tension, *V* is the box volume, and *ϕ*_0_ is the initial composition and the Lagrange multiplier *λ* ensures mass conservation. Since we treat the total volume *V* as a parameter there are four unknowns, *ϕ*_*in*_, *ϕ*_*out*_, *f* and *λ* in the above free-energy. Minimisation w.r.t these four unknowns leads to the following equations,

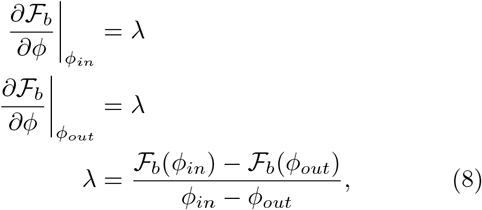

These three equations can be solved to yield the three unknown variables *ϕ*_*in*_, *ϕ*_*out*_, and *λ*. It should be noted that upon rearranging the three above equations, one arrives at the familiar common-tangent conditions: *µ*(*ϕ*_*in*_) = *µ*(*ϕ*_*out*_) and Π(*ϕ*_*in*_) = Π(*ϕ*_*out*_), where 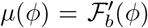 is the chemical potential and the osmotic pressure is similarly given by 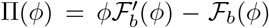. These two conditions ensure chemical and mechanical equilibrium, respectively.Thermal equilibrium is ensured as the Helmholtz free-energy is defined in a constant temperature ensemble. Once we know these, the solvent fraction can be found out from the fourth equation *∂* ℱ _*b*_ (*ϕ*)*/∂λ* = 0, which yields, 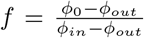. Note that these four equations are decoupled as first three equations do not involve the solvent fraction *f*. The situation is different for a spherical interface and that introduces a non-trivial coupling which we discuss in detail. Once these unknowns are determined, we are free to take the thermodynamic limit, which ensures that the effect of the interface term vanishes as *V*→ ∞. The equilibrium configuration is characterised by two coexisting phases with a planar interface as shown in the inset of Fig.(1) of the main manuscript. The interfacial tension between the coexisting phases has the form,

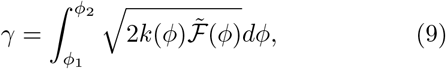

where 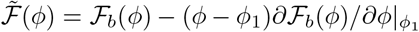 is the freeenergy after subtracting the common tangent, and *k*(*ϕ*) is the energetic cost associated with spatial variations of order parameter *ϕ* [**?**]. *k*(*ϕ*) has dimensions of ∼ *a*^2^, where *a* is the microscopic Kuhn length.

### B. The spherical interface

For a system with spherical interface (see inset of (b) of Figure 1 of the main manuscript), the Helmholtz freeenergy per unit volume of the droplet phase has the following form:

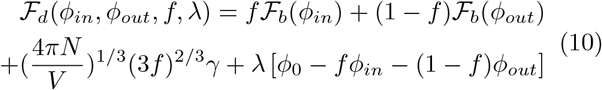

Minimising with respect to the four unknowns result in the following equations,

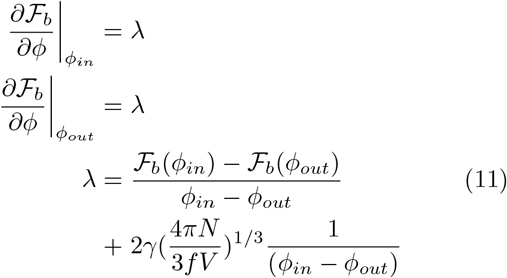

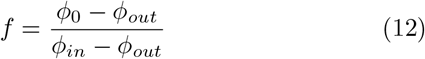

The first two equations imply the equality of chemical potentials: *µ*(*ϕ*_*in*_) = *µ*(*ϕ*_*out*_) and upon substitut-ing the value of *λ* from the third equation into the first and second one arrives at the second condition: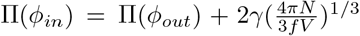. Now by substituting *f* from the last equation one ends up in the two equilibrium conditions expressed in terms of the coexistence densities *ϕ*_*in*_ and *ϕ*_*out*_ and they are solved numerically to yield the coexistence densities for a given box volume *V* and surface tension *γ*.

### C. The elastic free-energy

The total elastic free energy associated with accommodating a single solvent droplet of radius, *R*, inside the gel of mesh size, *R*_0_ is given by [14],

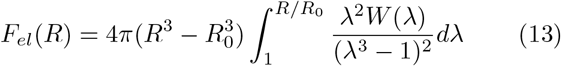

To incorporate the effects of the finite stretch-ability of the gel, we adopt Gent model [32, 33], where the elastic free energy density is of the form, 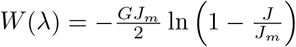, where 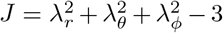, *J*_*m*_ ∼ 10^6^ is an upper-limit of the stretching and *G* is the shear modulus of the network gel. The shear modulus is related to the microscopic parameters via the relation, 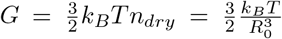, where *n* _*dry*_ is the cross-link density of the dry-gel and *R*_0_ is the mesh size of the dry gel [34]. The mesh size *R*_0_ is given by 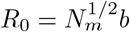, where *N*_*m*_ is the number of monomers along the backbone of the dry gel, between two cross-links (*N*_*m*_ = 16 in our subsequent calculations). The parameter *b* is the linear dimension of an effective polymeric monomer and following Tanaka [34] we take *b* ∼5*a*, where *a* is the Kuhn length or the smallest length-scale associated with the polymer. Due to the volume-preserving nature of the deformation, one has 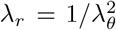 and *λ*_*ϕ*_ = *λ*_*θ*_ = *λ* [32] and the deformation obeys the following bound: 0 *< J/J*_*m*_ *<* 1.

The upper limit of the above integral signifies the droplet-gel interface and the lower limit signifies a region far away from the centre of the droplet where the stress fields have decayed and the gel there is completely unstressed. However, in a gel with N droplets of the the solvent the stress field associated with the presence of a single droplet does not fully decay when one encounters the surface of the next droplet. To consider this effect we assume that the droplets are of uniform size and are in the equilibrium state sit on a cubic lattice. In this situation the distance between the centres of two neighbouring droplets is Δ = *L/N* ^1*/*3^ (see inset of Figure (5)) and thus the upper-cutoff distance which would be associated with the lower limit in the elastic-energy integral is *R*_*m*_ = Δ −*R*. If we consider the un-deformed distances then *R*_*m*_ would correspond to a radial distance of 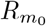, which would obey the following volume conserva-tion constraint, 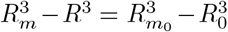 and this would lead to 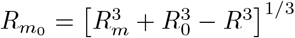 and thus the hoop-stretch or the lower limit of the elastic free-energy integral would be 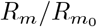. Thus the final form for the elastic energy per unit volume is,

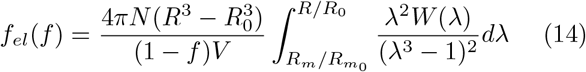

where the normalisation by the volume of the gel inside the box is evident and the transformation of the droplet radius in the elastic free-energy integral to the solvent fraction, *f* would occur via the relation 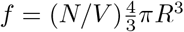. Thus the elastic free-energy per unit volume is thus expresses in terms of the solvent volume fraction, *f*. The upper panel of Fig. (6) shows the upper and the lower limits of the elastic energy integral expressed as a function of the solvent fraction *f*.

The primary set of parameters on which the thermodynamic phase of the system depends are: the surface energy *γ*, the shear modulus of the gel, *G*, which again depends on the mesh-size of the gel, *R*_0_. The presence or absence of the dispersed micro-droplet phase depends on the relative weights of the elastic and the surface-energy interactions [29]. In the limit where one has a single macroscopic solvent droplet of radius *r* → ∞ inside the gel the associated elastic energy per unit volume can be written in the form, *f*_*el*_(*r*) = *αG*, where *α* ∼1.5 is a dimensionless constant. In the limit where there are micro-droplets of solvent dispersed inside the gel, the surface tension becomes important. The surface energy per unit volume of the droplets is given by 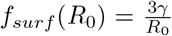. The ratio of the surface and the elastic energies per unit volume is given by,

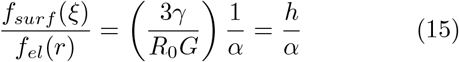

where, the dimensionless elasto-capillary number is given 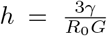. If *h < α*, then the thermodynamic stable state is that of dispersed micro-droplets in the gel, while if *h > α*, the stable phase is one with a single macroscopic droplet. In the subsequent calculations we choose a value of *γ* such that *h* ∼*α* and we go on to see whether we indeed observe a dispersed droplet phase in a more detailed microscopic mean-field theory calculations. The value of *γ* is set to 1/600, unless in the set calculations where the characteristics of the droplet phase is investigated by varying *γ* whule keeping *G* fixed.

Thus the total free-energy of the dispersed droplet phase in the background gel-matrix has the following form, when every term is expressed as a function of the solvent volume fraction, *f*,

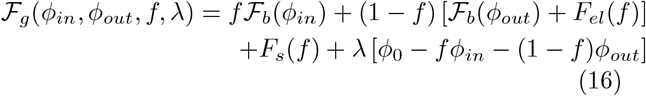

Upon minimising the above free-energy w.r.t the four unknowns one has the following equations,

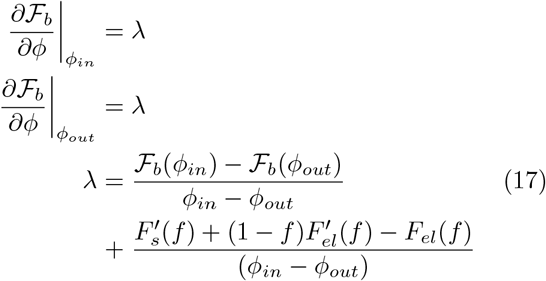

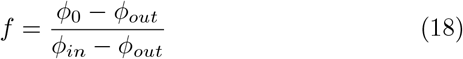

Again, by substituting the expression for *λ* into the first two equations and by identifying that 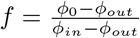, the above equations can be recast into two equilibrium conditions which impose chemical and mechanical equilibrium, respectively,

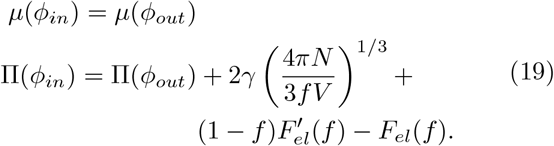

These can be solved numerically, via the parallel tangent construction, for the two coexistence densities provides one inputs the values of the surface tension and the box volume and the composition *ϕ*_0_. Figure (5) shows a schematic of the parallel tangent construction on the free-energy, ℱ_*b*_(*ϕ*), which has only a single minimum owing to the absence of translational entropy of the gel component. The point marked *ϕ*_0_ denotes the initial unstable phase composition, which ultimately splits into two coexisting phases at *ϕ*_*in*_ and *ϕ*_*out*_. As a result of the functional form of ℱ_*b*_(*ϕ*), *ϕ*_*in*_ is located at the unstable part of the free-energy surface. In spite of this, the system is maintained in mechanical equilibrium via the elastic energy residing in the gel-matrix.

The numerical solution of these two equations proceeds in the following manner: we always ensure that the chemical-equilibrium is maintained by chosing a pair of points *ϕ*_*in*_ and *ϕ*_*out*_, such that the local tangents are parallel (*µ*(*ϕ*_*in*_) = *µ*(*ϕ*_*out*_)). Since the value of the composition, *ϕ*_0_, is an input, the value of the solvent fraction, *f*, is readily computed. One then plots the three terms in the appearing in the mechanical equilibrium condition as a function of the solvent volume fraction, *f*, and search for the intersection of these curves (see the lower panel of Fig. (6)). This procedure is then repeated for various number of droplets.

After each of these four unknown variables, *ϕ*_*in*_,*ϕ*_*out*_,*f*, and *λ* have been determined from the computation associated with each droplet number, *N*, the equilibrium values are substituted back into the original free-energy expression (see Eq. (16)), which we set out to minimise, we get a new free-energy, 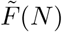, which is a function of the number of droplets *N*. We explore the properties of 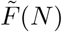 to investigate its shape and whether it admits a single or a multiple droplet minimum.

